# Gene regulatory dynamics during the development of a paleopteran insect, the mayfly *Cloeon dipterum*

**DOI:** 10.1101/2024.05.14.594094

**Authors:** Joan Pallarès-Albanell, Laia Ortega-Flores, Tòt Senar-Serra, Antoni Ruiz, Josep F. Abril, Maria Rossello, Isabel Almudi

**Affiliations:** Department of Genetics, Microbiology and Statistics, Universitat de Barcelona, Diagonal 643, 08028, Barcelona, Spain; Institut de Recerca de la Biodiversitat (IRBio), Universitat de Barcelona, Diagonal, 643, 08028, Barcelona, Spain; Institute of Biomedicine of Universitat de Barcelona (IBUB), Universitat de Barcelona, Diagonal, 643, 08028, Barcelona, Spain

**Author notes:** Equally contributed (Joan Pallarès-Albanell, Laia Ortega-Flores).

**Keywords:** Gene regulation, Paleoptera, mayflies, insects, embryogenesis, ATAC-seq

## Abstract

The evolution of insects has been marked by the appearance of key body plan innovations and novel organs that promoted the outstanding ability of this lineage to adapt to new habitats, boosting the most successful radiation in animals. To understand the origin and evolution of these new structures, it is essential to investigate which are the genes and gene regulatory networks participating during the embryonic development of insects. Great efforts have been made to fully understand, from a gene expression and gene regulation point of view, the development of holometabolous insects, in particular *Drosophila melanogaster*, with the generation of numerous functional genomics resources and databases. Conversely, how hemimetabolous insects develop, and which are the dynamics of gene expression and gene regulation that control their embryogenesis, are still poorly characterized. Therefore, to provide a new platform to study gene regulation in insects, we generated ATAC-seq (Assay for transposase-Accessible Chromatin using sequencing) for the first time during the development of the mayfly *Cloeon dipterum.* This new available resource will allow to better understand the dynamics of gene regulation during hemimetabolan embryogenesis, since *C. dipterum* belongs to the paleopteran order of Ephemeroptera, the sister group to all other winged insects. These new datasets include six different time points of its embryonic development and identify accessible chromatin regions corresponding to both general and stage-specific promoters and enhancers. With these comprehensive datasets, we characterised pronounced changes in accessible chromatin between stages 8 and 10 of embryonic development, which correspond to the transition from the last stages of segmentation to organogenesis and appendage differentiation. The application of ATAC-seq in mayflies has contributed to identify the epigenetic mechanisms responsible for embryonic development in hemimetabolous insects and it will provide a fundamental resource to understand the evolution of gene regulation in winged insects.

## INTRODUCTION

Insects are the most numerous and diverse lineage of animals on the planet (Misof et al. 2014). This huge radiation has been possible due to their extraordinary capabilities to adapt to distinct environments and the myriad of life history traits evolved within this animal class, which resulted in more than thirty extant orders distributed worldwide (Misof et al. 2014; D. Grimaldi & M. S. Engel 2007).

This diversity of forms and lifestyles is the result of changes in the gene regulatory networks (GRNs) controlling the embryonic and postembryonic development of this clade (Carroll 1998; Molina-Gil et al. 2023). In recent decades, the importance of regulatory information responsible for the location and time in which genes and GRNs are functioning, has been widely recognised (Furlong and Levine 2018; Leyhr et al. 2022; Gompel et al. 2005; Andrikou and Arnone 2015). These so called *cis* regulatory elements (CREs) are major players of morphological evolution not only in insects, but also in other animal lineages (Guerreiro et al. 2013; Leyhr et al. 2022). Still, some changes in trans, - in the coding sequences of transcription factors (TFs) and/or signalling molecules-have been also shown to participate in the diversity of phenotypes observed in animals (Galant and Carroll 2002; Santos et al. 2017). Thus, both changes in coding regions and CREs are key to explain the wide range of insect (and animal) morphologies. Comparative genomics projects (Crowley et al. 2023; Formenti et al. 2022; Molik 2022) promoted a better understanding of changes in trans elements, due to their higher degree of conservation between species. By contrast, CREs are more difficult to identify by homology, since they are usually less pleiotropic and for this, they tend to accumulate more changes in their sequences that impede their proper characterisation by sequence similarity. This is even more manifest in insects, with increased rates of genome evolution when comparing with vertebrate lineages (Zdobnov and Bork 2007). In addition, comparative transcriptomics provide information about the spatio-temporal dynamics of gene expression between different lineages (Mantica et al. 2024; Rodríguez-Montes et al. 2023; Levin et al. 2016), which may reflect to some degree differences in GRNs. The advent of new functional genomics approaches based on chromatin accessibility, such as, first, formaldehyde-assisted isolation of regulatory elements (FAIRE-seq (J. M. Simon et al. 2012)) and the more recent Assay for Transposase-Accessible Chromatin (ATAC-seq (Buenrostro et al. 2013)), allow the identification of open chromatin regions that can be assigned as CREs, such as enhancers, promoters and insulators. The advantages of these methods, in comparison to other functional genomics assays used before (e.g. ChIP-seq), are the low input of starting material, the high-throughput protocol without expensive reagents or antibodies and that they are not excessively laborious, among others. Therefore, much more detailed and comprehensive characterisations of CREs have been possible and several FAIRE-seq and ATAC-seq datasets have been generated during the last decade to address distinct questions related to insect development and evolution. Nonetheless, these works have been mostly done using few established model species, such as the fruitfly *Drosophila melanogaster* (Diptera), the red flour beetle *Tribolium castaneum* (Coleoptera), some butterfly and moth species (Lepidoptera) or certain species of ants and bees (Hymenoptera) (Lowe et al. 2022; Wang et al. 2020; Zhao et al. 2020), all of which belong to a single monophyletic group of insects, the Holometabola (Fig. 1A)(Schmidt-Ott and Lynch 2016). By contrast, the remaining seventeen orders of winged insects, the Hemimetabola (Fig. 1A), continue unexplored, with only adults and one embryonic FAIRE-seq datasets available (Fernández et al. 2020; Richard et al. 2017) to our knowledge.

**Figure 1.**
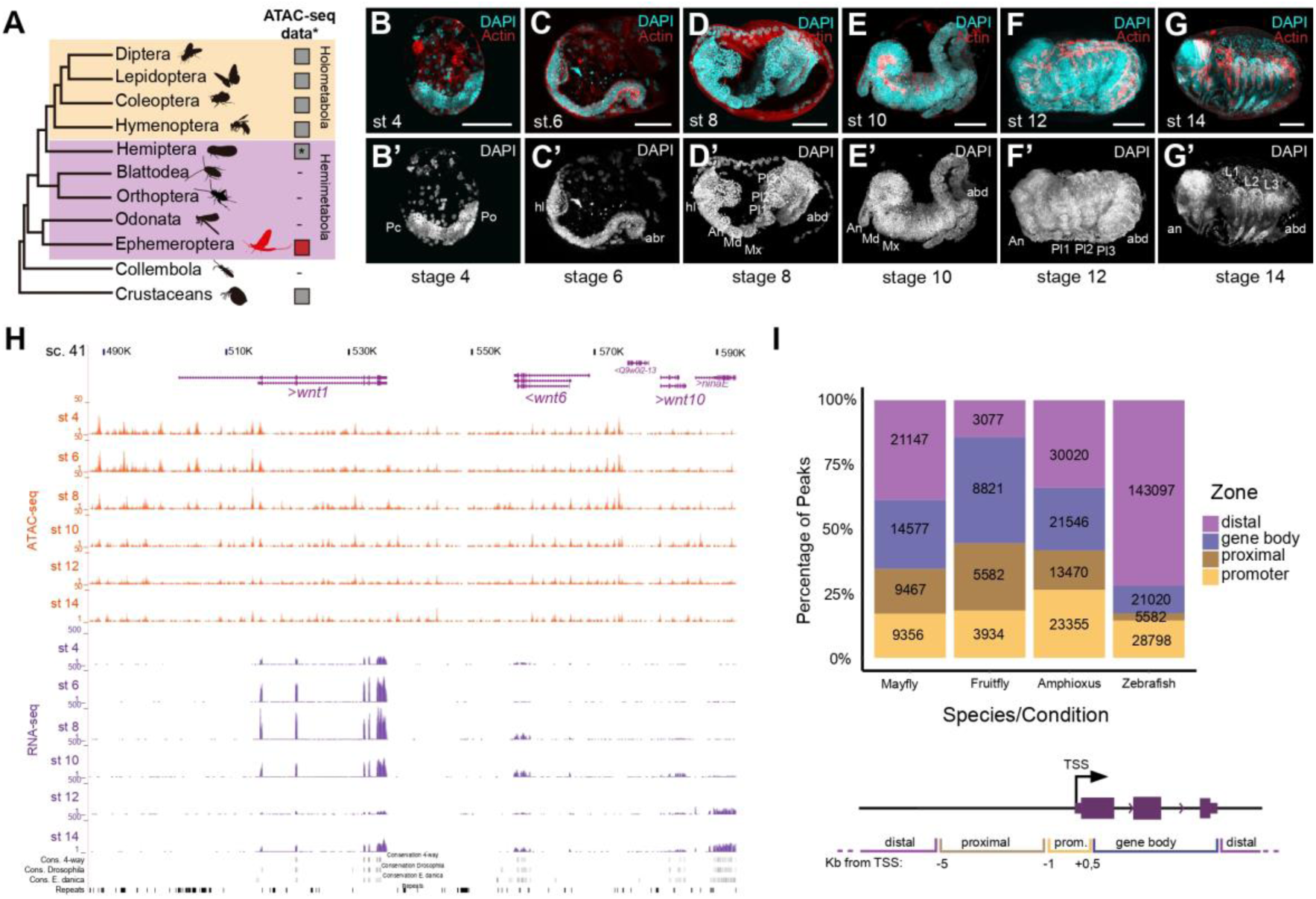
Open Chromatin profiles in *C. dipterum* embryogenesis. **(A)** Simplified insect phylogeny. Grey squares highlight the availability of ATAC-seq datasets. Asterisk shows lineages in which there is no ATAC-seq information but FAIRE-seq material has been generated. **(B-G)** Embryonic stages used in this study stained with Phalloidin-AlexaFlour-488 (Red) and DAPI (cyan). (B-B’) Stage 4 embryo (St 4): germ band initial elongation. (C-C’) St 6: S-shaped embryo: anatrepsis II. (D-D’) St 8: Segmentation of the embryo. (E-E’) St 10 embryo: revolution or katatrepsis. (F-F’) St 12: initial dorsalization. (G-G’) St 14 embryo: dorsal closure completed. **(H)** Snapshot of the UCSC genome browser showing *wnt1, wnt6, wnt10* cluster and the ATAC-seq tracks in this study and and RNA-seq tracks generated in this study and in (Almudi et al. 2020). **(I)** Percentage of ATAC-seq APREs distributed across *C. dipterum*, *D. melanogaster, B. lanceolatum* and *D. rerio* genomes. Scale bars: 50 um.

Both hemimetabolous and hemimetabolous orders shared common phases during their development (Sander 1976; Patel 1994). Indeed, these main events during embryogenesis are also present in non-insect arthropods (Zrzavý and Štys 1997). The early stages of insect development initiate with several rounds of nuclear divisions that migrate to the periphery to form the blastoderm. The blastoderm gives rise to different types of germ band (short, intermediate or long) which are subsequently segmented during the following embryonic stages (Davis and Patel 2002). Then, the segmented embryo undergoes a process of differentiation in which organogenesis and the final development of the juvenile structures take place (Sander 1976).

The mayfly *Cloeon dipterum,* a recently established laboratory system (Almudi et al. 2020; 2019), is in a privileged position to improve the phylogenetic diversity of insect functional genomics resources. Mayflies or Ephemeroptera belong to the Paleoptera group of winged insects, together with Odonata (dragonflies and damselflies)(Fig. 1A) (S. Simon, Blanke, and Meusemann 2018). They are the sister group to all other winged insects and thus, due to this position in the insect phylogenetic tree, they are key to address fundamental questions related to insect physiology, ecology, development and evolution.

Here, we performed ATAC-seq experiments at six different developmental stages, -including some of the hallmark stages of insect embryogenesis mentioned above-, in *C. dipterum* embryos. Our identified accessible chromatin regions provide an exhaustive collection of putative promoters and enhancers along embryogenesis of this hemimetabolan insect. Moreover, by studying the temporal dynamics of these elements, we showed wholesale changes in chromatin accessibility during the transition between the last stages of segmentation and the start of organogenesis and appendage differentiation. Finally, we facilitate the access to these comprehensive datasets through a dedicated web browser (https://genome-euro.ucsc.edu/s/mayfly/Clodip), providing a key resource available for the entire community to understand the evolution of gene regulation during the development of winged insects.

## RESULTS AND DISCUSSION

### Open Chromatin profiles in *C. dipterum* embryogenesis

To investigate dynamics of gene regulation during the embryogenesis of mayflies, we performed ATAC-seq assays for six different developmental stages: stage (st) 4 (germ band elongation, Fig 1B, B’), st 6 (S-shaped embryo: anatrepsis II Fig. 1C, C’), st 8 (segmentation of the embryo, Fig. 1D, D’), st 10 (revolution: katatrepsis, Fig. 1E, E’), st 12 (start of dorsal closure, Fig. 1F, F’), st 14 (dorsal closure complete, Fig. 1G, G’)(Tojo and Machida 1997). Sequences resulting from these experiments were mapped against the *C. dipterum* reference genome assembly (CLODIP2 (Almudi et al. 2020), see methods and Fig. S1) to obtain a non-redundant collection of open chromatin regions throughout the genome that we termed APREs (Accessible Putative Regulatory Elements) (Fig. 1H). After normalization (see methods) we identified a total of 54,547 APREs across the six developmental stages. Of them, 45,649 APREs did not show changes in accessibility in our clustering analyses of the different developmental samples we assayed (i.e. they remained constitutively open or close across these stages) while 8,898 APREs were dynamic and changed their accessibility across developmental stages (Fig. S2 and Table S1).

We next aimed at defining the genomic distributions of the APREs relative to genes and gene annotations. For this, we calculated the proportions of APREs at the “promoter” (i. e. APREs in the immediate vicinity of the annotated transcription start sites (TSSs)), at “proximal regions”, spanning up to 5 Kb upstream the TSSs, at “gene bodies” (located between the end of the promoter and the termination site of the gene) and “distal” regions, which comprised genomic regions that do not fall in the previous categories (Fig. 1I, Table S2). We found that both non-dynamic and dynamic APREs were distributed in similar proportions (Fig. S3) and only detected a slight increase in non-dynamic APREs located in promoters with respect to dynamic APREs in promoters (∼18% versus ∼13%) and an even slighter difference between non-dynamic and dynamic APREs in gene bodies (26% and 29%, respectively, Fig. S3). This proportion of APRE distribution was similar to the distribution of APREs in other invertebrate genomes, such as the chordate amphioxus (*Branchiostoma lanceolatum*) (Marlétaz et al. 2018). These two invertebrate species, *C. dipterum* and *B. lanceolatum,* notably diverged from the distribution found in some vertebrate species, which showed much larger proportion of distal APREs due to the relevance of distal regulation in these vertebrate genomes and the impact of the different rounds of whole genome duplications (Marlétaz et al. 2018). By contrast, when we compared the APRE distribution of *C. dipterum* with the distribution of accessible chromatin in *D. melanogaster,* we observed that fruitflies had a much lower proportion of distal APREs (i.e. a third of the corresponding fractions in *C. dipterum* and *B. lanceolatum*), with a higher proportion of APREs located in gene bodies and proximal regions (Fig. 1I)(Bozek et al. 2019). These differences between *C. dipterum* and *D. melanogaster* were most likely due to the higher compaction of the 120 Mb *D. melanogaster* euchromatic genome (Adams et al. 2000), where most of the genomic regulatory blocks ancestral to animals and their associated long range regulatory interactions have been dismantled (Irimia et al. 2012).

### ATAC-seq revealed a temporally regulated chromatin profile in the mayfly genome

In order to study changes in chromatin accessibility throughout different samples (i.e. developmental timepoints), we analysed differential APRE activity between consecutive developmental stages. These results showed relatively modest changes between successive timepoints (i.e. 164 APREs at the St 4 to St 6 transition or 365 at St 10 to St 12, Fig. S4, Table S3), with the notable exception of the transition between St 8 and St 10, when more than four thousand APREs (4568) changed their accessibility, from 7,5 to 27 times more than at the other timepoints (Fig. S4). From these, the vast majority (3118) corresponded to APREs that increase accessibility during this St 8-St 10 phase. This large amount of differentially active APREs between these two stages may indicated a major regulatory turnover between an early and late regulatory state during mayfly embryogenesis. In fact, this major shift was also evident when we performed a PCA analysis and clustering of the ATAC-seq datasets, which formed two very distinctive clusters: samples from stages 4, 6 and 8 and samples from 10, 12 and 14 (Fig. S4).

To further explore the dynamics of chromatin accessibility across the selected developmental timepoints, we performed a temporal soft-clustering analysis using Mfuzz (Kumar and Futschik 2007)(see methods and Table S4). Among the different temporal clusters obtained (Fig. S5), we focused on clusters of APREs whose accessibility peaked at a single embryonic stages (e.g. cluster 8 and 16 for st 4, cluster 29 and 21 for st 6, cluster 28, 5, 6, 11, 13, 14 and 26 for st 8, cluster 25, 15 and 18 for st 10, cluster 27 for st 12 and cluster 4 and 2 for st 14; Fig 2A and Fig. S5). We then associated these APREs to their putative target genes and analysed the Gene Ontology (GO) enriched terms for each of these stage-specific clusters, using *D. melanogaster* orthologs (Almudi et al. 2020), since it is the closest organism with functional annotation available (see methods and Table S5). This GO term enrichment analysis exhibited categories highly connected with each of the embryonic stages in which the accessibility of the chromatin was higher (Fig. 2A and Table S5). For instance, cluster 8 (st 4) and cluster 29 (st 6) contained APREs that were associated to genes involved in cell adhesion, planar cell polarity, DNA biosynthetic process and negative regulation of cell differentiation, which are characteristic processes of early embryogenesis in insects (Münster et al. 2019; Brantley and Di Talia 2021). On the other hand, GO terms enriched in clusters corresponding to later stages of embryogenesis (e.g. cl 25, cl 27 and cl 4, Fig 2A and Table S5) revealed processes related to organogenesis, such as axon guidance or regulation of developmental process (Gillott 2005).

**Figure 2.**
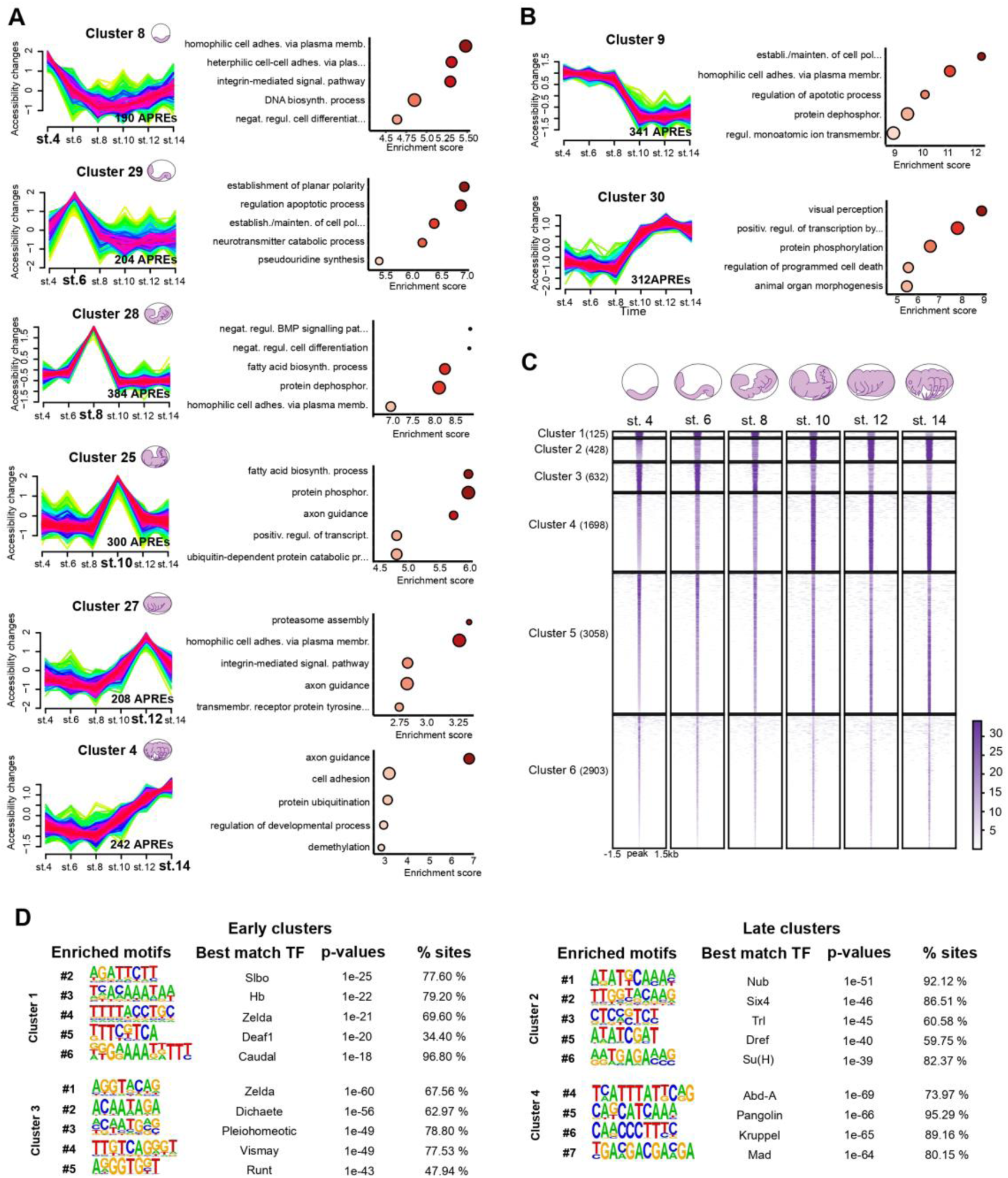
ATAC-seq revealed a temporally regulated chromatin profile in the mayfly genome. **(A)** Mfuzz clusters obtained for the 8,898 dynamic APREs obtained through ATAC-seq experiments in the six selected developmental stages representing stage-specific APRE activity and their associated GO enriched terms. **(B)** Mfuzz clusters representing “early embryogenesis” and “late embryogenesis” APRE activity and their associated GO enriched terms. **(C)** Heatmaps of the 8,898 dynamic APREs clustered using k-means clustering. Six clusters were obtained, with four of them showing a clear dynamic behavior: cluster 1 (n=125) and cluster 3 (n=632) or early activity, cluster 2 (n=428) and cluster 4 (n = 1698) or late activity. **(D)** Motif enrichment analysis of the early (cluster 1, cluster 3) and late (cluster 2, cluster 4) active clusters. Five or four representative motifs of the top-10 were chosen. Motif logos are represented with their position in the top-10, the TF names, the enrichment p-values and the percentage of sites showing the motif.

Besides these stage-specific clusters, we also found several clusters that showed more prolonged activity patterns. In this manner, we identified clusters that recapitulated the major developmental shift we had previously observed between st 8 and st 10, with a set of “early embryogenesis” clusters (e.g. clusters 7, 9, 17, 19, 21 or 26) and another of “late embryogenesis“ones (clusters 3, 22, 23 or 3; Fig. 2B and Fig. S5). Accordingly, we identified enriched GO terms related to early development (e.g. establishment and maintenance of cell polarity or cell adhesion processes) and terms related to late embryogenesis (e.g. visual perception and animal organ morphogenesis), respectively (Fig. 2B and Table S5).

In addition, we also performed *k-means* hard clustering (see methods, Table S6) using the same set of dynamic APREs (Fig. 2C). This analysis was able to recover six clusters with differential dynamics of accessibility, although none of these clusters corresponded to APREs showing stage-specific accessibility. By contrast, we identified two groups of clusters, - cluster 1 and cluster 3, on one hand, and cluster 2 and cluster 4, on the other hand-, that contained APREs accessible during early stages of embryogenesis and late stages of embryogenesis, respectively (Fig. 2C and Table S6), mirroring the st 8-st 10 shift observed in the previous analyses. In order to better characterise these clusters, we carried out TF motif enrichment analysis using Homer software (see methods, Fig. 2D and Table S7). In agreement with previous work in other insects and the results of the GO term enrichment analysis from the Mfuzz clusters (Fig. 2A, B), for APREs in cluster 1 and cluster 3, we identified motifs whose best match were TFs involved in early embryogenesis. These include Hunchback (Hb), a gap gene involved in antero-posterior axis specification (Qian, Capovilla, and Pirrotta 1991), Caudal (Cad), which functions in germ band elongation (Schulz and Tautz 1995; Wu and Lengyel 1998), or Zelda (Zld), a zygotic genome activator that acts during early blastoderm development (Brennan et al. 2023; Liang et al. 2008) (Fig. 2D). By contrast, clusters 2 and 4, whose APREs were open in later stages of development, showed enrichment in motifs that correspond to TFs involved in different processes of organogenesis, such as Nubbin (Nub), a regulator of appendage morphogenesis (Turchyn et al. 2011), Six4, involved in the development of mesodermal structures (Clark et al. 2006), or Mothers against dpp (Mad), that mediates the response to the BMP pathway during the development of diverse insect organs (Sekelsky et al. 1995).

Overall, these analyses revealed two main phases during the embryogenesis of mayflies in which distinct set of regulatory regions are active (Fig. 2B, C) to control different sets of genes and regulatory networks involved in such early or late embryonic processes. These results were consistent with the mid-developmental transition previously described at transcriptomic level for some phyla, including insects (Levin et al. 2016).

### Chromatin accessibility to understand gene expression dynamics

Since ATAC-seq has been proven to be a powerful method to investigate the regulation of gene expression, we next addressed the relationship between chromatin accessibility and levels of gene expression (Starks et al. 2019). To do this, we measured the levels of gene expression at the same developmental stages of our ATAC-seq datasets (see methods Fig. S6 and Table S8). When focusing on genes associated to the 8,898 dynamic APREs, we first observed that the six stages clustered accordingly to the “early” and “late” embryonic phases that we identified in the ATAC-seq data. Transcriptomes from stages 4, 6 and 8 formed a cluster while transcriptomes from stages 10, 12 and 14 grouped together (Fig. 3A). Moreover, we detected some genes whose expression varied along the developmental timepoints we characterised: cluster 11 and cluster 7 decreased their expression as embryogenesis progressed, while genes from cluster 3, 5 or 9 increased their expression during embryonic development (Fig. 3A).

**Figure 3.**
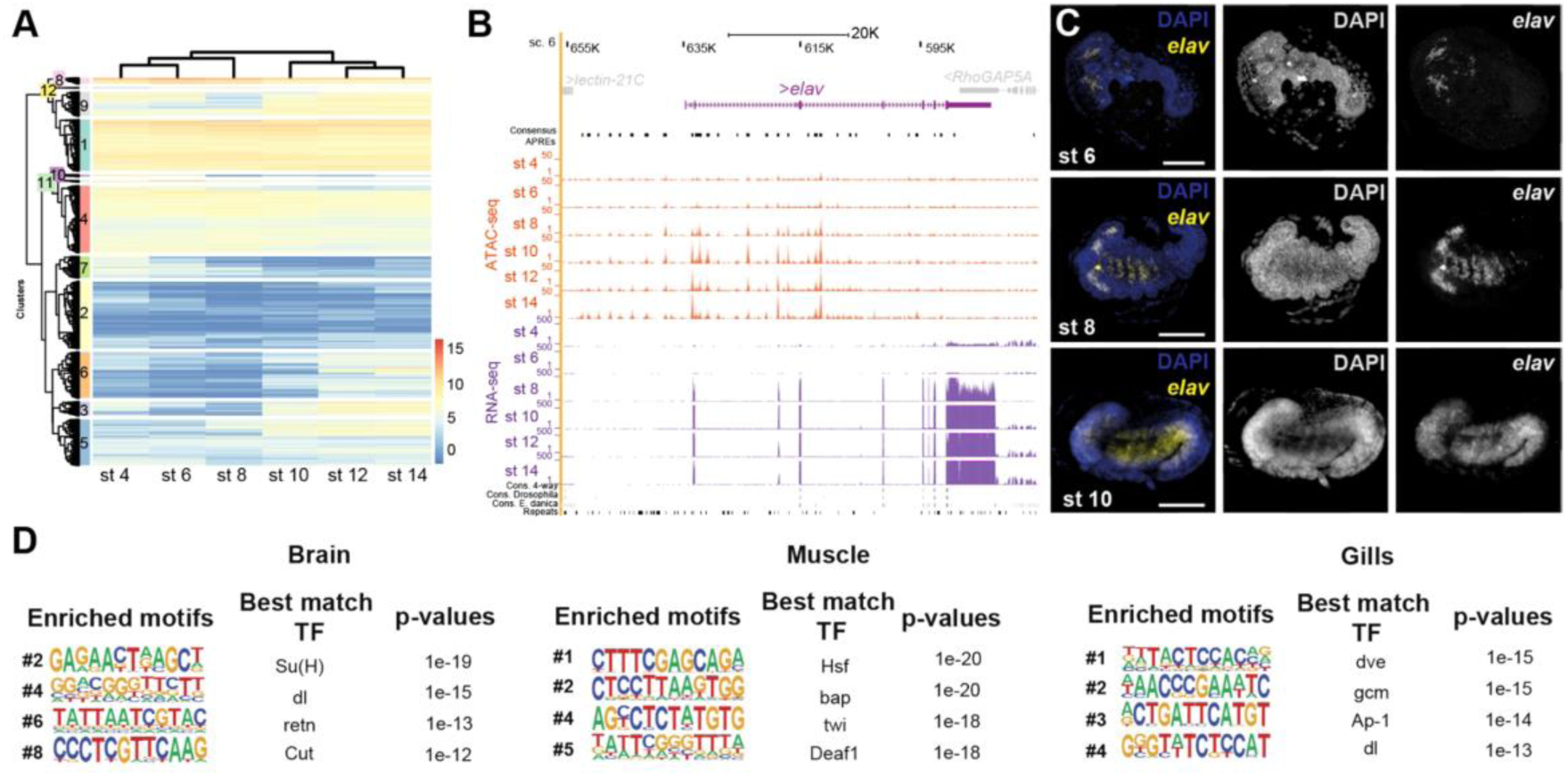
Chromatin accessibility to understand gene expression dynamics. **(A)** Heatmap showing expression levels of genes associated to dynamic ATAC-seq APRE in the selected embryonic stages. RNA-seq samples clustered according to embryonic stage progression. Secondary clustering showed 12 different gene clusters. **(B)** *elav* genomic regulatory landscape. APRE activity and gene expression increase as embryogenesis progresses. **(C)** HCR hybridization against the gene *elav* at st 6, st 8 and st 10 of embryogenesis. Nuclei were stained with DAPI (dark blue) and *elav* expression pattern is shown in yellow. Scale bars: 50 um. **(C)** Motif enrichment analysis of APREs associated to genes from some tissue-specific WGCNA modules identified in (Almudi et al. 2020). Four representative motifs of the top-10 were chosen. Motif logos are represented with their position in the top-10, the TF names and the enrichment p-values.

To further illustrate our results, we investigated the expression pattern of *embryonic lethal abnormal vision* (*elav*) that showed differential APRE accessibility and differential gene expression along embryonic stages (Fig. 3B). *Elav* is a RNA binding protein involved in axon guidance, synapse formation and development and maintenance of neurons (Robinow and White 1988). We performed Hybridization Chain Reaction (HCRs) assays (Bruce et al. 2021) in st 6, st 8 and st 10 embryos to characterise the spatial expression of this gene in these stages in which we observed shifts in chromatin accessibility and expression levels. While embryos at st 6 showed reduced expression of *elav* in some cells in the cephalic region (Fig. 3C), at st 8 *elav* exhibited a broader expression domain in head domains and in the most anterior thoracic segments. At st 10, these neural territories of *elav* expression expanded and elongated to the abdominal segments (Fig. 3C). As expected, we also observed an increase in the accessibility of chromatin in the locus, especially upstream of the Transcription Start Site (TSS, Fig. 3B).

Previous analysis of co-regulated gene expression across several tissues, using Weighted Gene Correlation Network Analysis (WGCNA), revealed modules of genes specifically expressed in particular adult and nymphal tissues (Almudi et al. 2020). We examined whether chromatin accessibility information from embryos correlated with these modules. We characterised enriched motifs in APREs associated to the genes contained in each of these WGCNA modules and found distinctive enriched binding motifs for some of them (see methods and Table S9). For example, the brain module was enriched in neural motifs, such as Suppressor of Hairy (Su(H)), retained (retn) or Cut (Ct), which are TFs involved in neural or glial development (Grueber and Jan 2004; Shandala et al. 1999), the muscle module had motifs for bagpipe (bap) or twist (twi), required for mesodermal development (Castanon et al. 2001; Cripps and Olson 2002) and specification or the gills module with enrichment in motifs such as defective proventriculus (dve) involved in epithelium patterning or glial cells missing (gmc), which could have a role in the determination of some of the numerous neural cells that we previously identified in these abdominal structures (Almudi et al. 2020; Hosoya et al. 1995) (Fig. 3D, and Table S9). These results suggest that some of the Gene Regulatory Networks involved in the development of nymphal tissues and organs are already functioning during embryogenesis and their regulatory signatures can be detected in our ATAC-seq datasets. Thus, our results can also provide important insights into the regulatory logic of the adult body plan, and therefore also constitute a valuable resource for adult insect biology.

Overall, our ATAC-seq datasets provide a comprehensive resource to help uncovering developmental diversity of insects, since it represents the first publicly available genome-wide collection of putative regulatory elements across embryogenesis in a hemimetabolous lineage using ATAC-seq approaches. Thus, the key phylogenetic position of Ephemeroptera, together with the extensive chromatin accessibility information made available here, will open new venues to address longstanding questions in the fields of developmental and evolutionary biology and comparative genomics.

## MATERIALS & METHODS

### Culture maintenance, embryo collection and fixation

Samples were obtained from a *Cloeon dipterum* culture maintained in the laboratory as previously described in (Almudi et al. 2019). Gravid females fertilised different days were collected and dissected to obtain embryos at selected developmental stages: st 4, st 6, st 8, st 10, st 12, st 14. After opening the abdomen of these gravid females, embryos were collected to perform ATAC-seq or RNA-seq procedures and a small subset was collected apart and fixed with 4% Formaldehyde for 1 hour at r.t. to confirm the developmental stage. After 3 x 5’ washes with PBS, these fixed embryos were stained with Phalloidin Alexa Fluor™ 488 (A12379) and DAPI to visualise actin filaments and nuclei, respectively. Images were acquired using a Zeiss LSM 880 confocal and were processed with Fiji (Schindelin et al. 2012).

### HCR hybridization

HCR hybridization followed a modified version of the Molecular Instruments (Los Angeles, CA, USA) HCR v.3 protocol (Bruce et al. 2021). HCR probe was designed to evade non-specific binding using an open-source probe design program (Kuehn et al. 2022). Briefly, embryos stored in ethanol were rehydrated in stepwise 75/50/25% ethanol in PBTw 0.1%. After 3 x 5’ washes in PBTw 0.1%, embryos were permeabilized in Detergent Solution (1.0% SDS, 0.5% Tween, 50.0 mM Tris-HCl (pH 7.5), 1.0 mM EDTA (pH 8.0), and 150.0 mM NaCl) for 30’ at room temperature (RT), kept in pre-warmed Probe Hybridization Buffer (Molecular Instruments) for 30’ at 37 °C, and incubated in Probe Solution (4 nM of probe in Probe Hybridation Buffer) overnight at 37 °C. After 4 x 15’ washes in pre-heated wash buffer (Molecular Instruments) at 37 °C and 2 x 5’ washes in 5x SSCTw 0.1% at RT, they were kept in pre-equilibrated Amplification Buffer (Molecular Instruments) for 30’ at RT and incubated in hairpin solution (60 nM of each hairpin h1 and h2 (Molecular Instruments) separately in pre-equilibrated Amplification Buffer, heated at 95 °C for 90 s and cooled down for 30’ overnight in the dark at RT. Following 5 x 20’ washes in 5x SSCTw 0.1% and 1 × 10’ wash in PBTw 0.1 % pH 7.4 in dark at RT, embryos were mounted in Prolongue™ Gold with DAPI (P36941, Invitrogen). Images were acquired using a Zeiss LSM 880 confocal and were processed with Fiji (Schindelin et al. 2012).

### RNA-seq sequencing and assembly

Three RNA-seq datasets (including replicates) of st 8 and st 12 embryos were generated using the Illumina technology. Samples were processed immediately after dissection and RNA was extracted using RNeasy Mini Kit (Qiagen) following manufacturers’ instructions. Paired-end libraries were generated using Illumina (Novaseq6000) 2×50bp. After quality control, the obtained reads were aligned using the STAR aligner. Initially, a genome index was created using the CLODIP2 reference genome (GCA_902829235.1) with the genomeGenerate mode of STAR. Subsequent alignment of reads to this index was performed using the alignReads mode. Gene expression levels were quantified utilizing the quantMode GeneCounts option within STAR.

### ATAC-seq and library preparation

ATAC-seq or assay for transposase-accessible chromatin by sequencing protocol was optimised during these experiments to use on mayflies from (Buenrostro et al. 2015). Briefly, embryos were homogenised in lysis buffer (10 mM Tris–HCl pH 7.4, 10 mM NaCl, 3 mM MgCl2, 0.1% NP-40) to obtain approximately 70000 individual nuclei. After removing lysis buffer, transposition reaction (1.25 μl of Tn5 enzyme in 10 mM Tris–HCl pH 8.0, 5 mM MgCl_2_, 10% w/v dimethylformamide) was performed for 30 min at 37 °C and the resulting fragments are purified using MinElute PCR Purification Kit (Qiagen). qPCR was performed to determine the optimal number of cycles necessary for each library. A unique pair of primers were assigned to each sample (Table S10) and 12 libraries were prepared corresponding to two biological replicates of the six selected developmental time points using a PCR. Libraries were purified using the MinElute PCR Purification Kit (Qiagen). DNA concentration in each sample was calculated with Invitrogen™ Qubit™ 4 Fluorometer using The Qubit 1X dsDNA HS Assay Kit.

### ATAC-seq mapping and peak (APRE) calling

For peak (APRE) calling, we followed the pipeline outlined in https://github.com/alexgilgal/Thesis_methods/tree/main/ATAC-seq%20analysis, implementing minor modifications as detailed in our project’s GitHub repository (https://github.com/mayflylab/Cdip-RegEmb/tree/main). We employed the ATAC_pipe.pl script for mapping reads, utilizing Bowtie2 (Langmead and Salzberg 2012) to align the reads to the CLODIP2 reference genome (GCA_902829235.1). Following alignment, the resulting BAM files were filtered based on a quality threshold of 10 and a minimum fragment length of 130 bp (Fig. S1).

The processed files were then subjected to peak analysis using the idr_ATAC_script.sh script, which executes peak calling with MACS2 (Zhang et al. 2008) to generate two sets of peaks: conservative peaks, indicating high-confidence regions across biological replicates, and optimal peaks, denoting reproducible events that consider read sampling variability, derived from pseudo-replicates. Subsequent IDR (Li et al. 2011) analysis was performed on both peak sets. Peak statistics—including the number of peaks and rescue ratios—were calculated and documented in a summary file.

### APRE classification and gene assignment

APREs are classified and associated with genes based on their proximity to the transcription start sites (TSS). TSSs are defined using the get_TSS.py script (https://github.com/m-rossello/GeneRegLocator/). To classify APREs and link them to genes, we use a custom-made script named make_table_from_zones.py (https://github.com/m-rossello/GeneRegLocator/). This script is designed to delineate regulatory zones around TSSs and associate these zones with APREs from ATAC-seq data. It defines three types of regulatory zones: Promoters, located near the TSS, spanning 1000 bases upstream and 500 bases downstream. Proximal regions, positioned further from the TSS, extending 4000 bases upstream but not overlapping with promoters. Gene bodies, encompassing regions within the gene but excluding the promoter areas (Fig. 1I). Zones are non-overlapping on the same strand, although the same genomic position can exhibit different zones on each strand. Each APRE is associated with one or more genes if it overlaps by more than 70% with a gene zone. APREs not falling within promoter, proximal, or gene body regions are classified as distal and remain unassociated with any gene (Table S2).

### Open chromatin analysis

The counts obtained from consensus APREs were used for all subsequent analyses, following normalization. This normalization involves adjusting the count data using the TMM method (Robinson and Oshlack 2010) to account for differences in library size and composition. To further explore trends and variations across different biological stages, we aggregate the normalized counts by these stages, calculating mean values. Global APRE analysis categorizes APRE as either “open” or “closed” based on a threshold of 10 counts. APREs registering fewer than 10 counts are deemed closed. We define non-dynamic APREs as those that remain consistently open or closed across all examined stages. Conversely, dynamic APREs are characterized by their variability, changing between open and closed states across different stages or samples.

### Differential Chromatin Accessibility Analysis

After counts normalization by TMM (Robinson and Oshlack 2010) and sample exploration by PCA and clustering, differential chromatin accessibility analysis was performed in dynamic APREs. For this analysis, we utilize the limma-trend method (Law et al. 2014; Phipson et al. 2016). This approach is applied to the normalized count data of dynamic APREs to identify significant differences in chromatin accessibility between conditions. To visualize the results, we generate volcano plots using the EnhancedVolcano package (Blighe, Rana, and Lewis 2023).

### Mfuzz analysis

We conducted the Mfuzz cluster analysis using the mfuzz function from the R package Mfuzz (Kumar and Futschik 2007). Dynamic APREs with mean values computed by developmental stage were analysed. The optimal parameters were systematically determined, setting the fuzzifier value at m=1.5 and the number of clusters at 30.

### k-means analysis

We performed k-means hard clustering using the DeepTools package (Ramírez et al. 2016) to analyze dynamic APRE enrichments from ATAC-seq data. This analysis included computing genome region scores with the computeMatrix function. The generated matrices enabled the visualization of heatmaps, which provided insights into the distribution of dynamic APREs across various developmental stages. Additionally, we created profile plots to further explore the chromatin accessibility dynamics.

### Gene Ontology Enrichment Analysis

For our Gene Ontology (GO) enrichment analysis, GO annotations are transferred from the UniProt proteome (UP000494165) to the genes associated with the APREs. We perform statistical analysis using the topGO package (Alexa and Rahnenfuhrer 2023), which utilizes the elimination algorithm to identify significantly enriched GO terms within gene clusters. The BH false discovery rate correction method (Benjamini and Hochberg 1995) is employed to control multiple testing. The results are visualized using GO enrichment plots, created with the ggplot2 package.

### TFBM enrichment analysis

Transcription factor binding motif (TFBM) enrichment analysis is conducted using the findMotifsGenome.pl tool from the HOMER suite (Heinz et al. 2010). We designate the APREs of interest as the foreground and utilize the remainder of the consensus APREs as the background. The fragment size selected for motif discovery corresponds precisely to the regions of the APREs (-size given). This analysis includes a comparison against collected motifs specific to insects.

## Acknowledgements

We thank María José López Fernández, Juan J. Tena, Alejandro Gil and Alberto Pérez-Posada for advice on bioinformatics analyses and Ignacio Maeso and the rest of the members of the Almudi lab for fruitful discussions. This research was funded by the European Research Council (ERC) under the European Union’s Horizon Europe research and innovation programme (ERC-CoG2021-101043751 to I.A), by the Spanish Ministry of Science and Innovation (PID2020-116041GB-I00 to IA) and by Ministry of Science, Innovation and Universities and by “European Union NextGeneration EU/PRTR” (CNS2023-145403). M. R. holds a Margarita Salas fellowship, from Spanish Ministry of Science and Innovation and I.A. is the recipient of the “Subprograma Estatal de Incorporación 2017-2020: Beatriz Galindo” (BG20--00115) from the Spanish Ministry of Universities.

## Author contributions

J.P., L.O-F and M.R. performed most analyses and generated most figures and tables. T.S and A.R performed spatial embryo fixations and HCR assays. L.O-F performed ATAC-seq experiments and generated the libraries. J.F.A and J.P. set up UCSC browser. I.A. and M.R. coordinated the project. I.A. obtained funding and wrote the main text with inputs from all authors.

## Data availability

The datasets generated and analysed during the current study are available in the Sequence Read Archive (SRA) repository. The ATAC-seq dataset, which includes samples from six developmental stages, can be accessed via SRA accession numbers SAMN40277609 through SAMN40277620. Additionally, RNA-seq data for three developmental stages are stored under SRA accession numbers SAMN40990690, SAMN40990692, and SAMN40990693. All datasets are grouped under the project accession number PRJNA1084266. Datasets are also available as a UCSC track hub: https://genome-euro.ucsc.edu/s/mayfly/Clodip. All code used for analysis is available on GitHub at https://github.com/mayflylab/Cdip-RegEmb/

## SUPPLEMENTARY FIGURES

**Figure S1.**
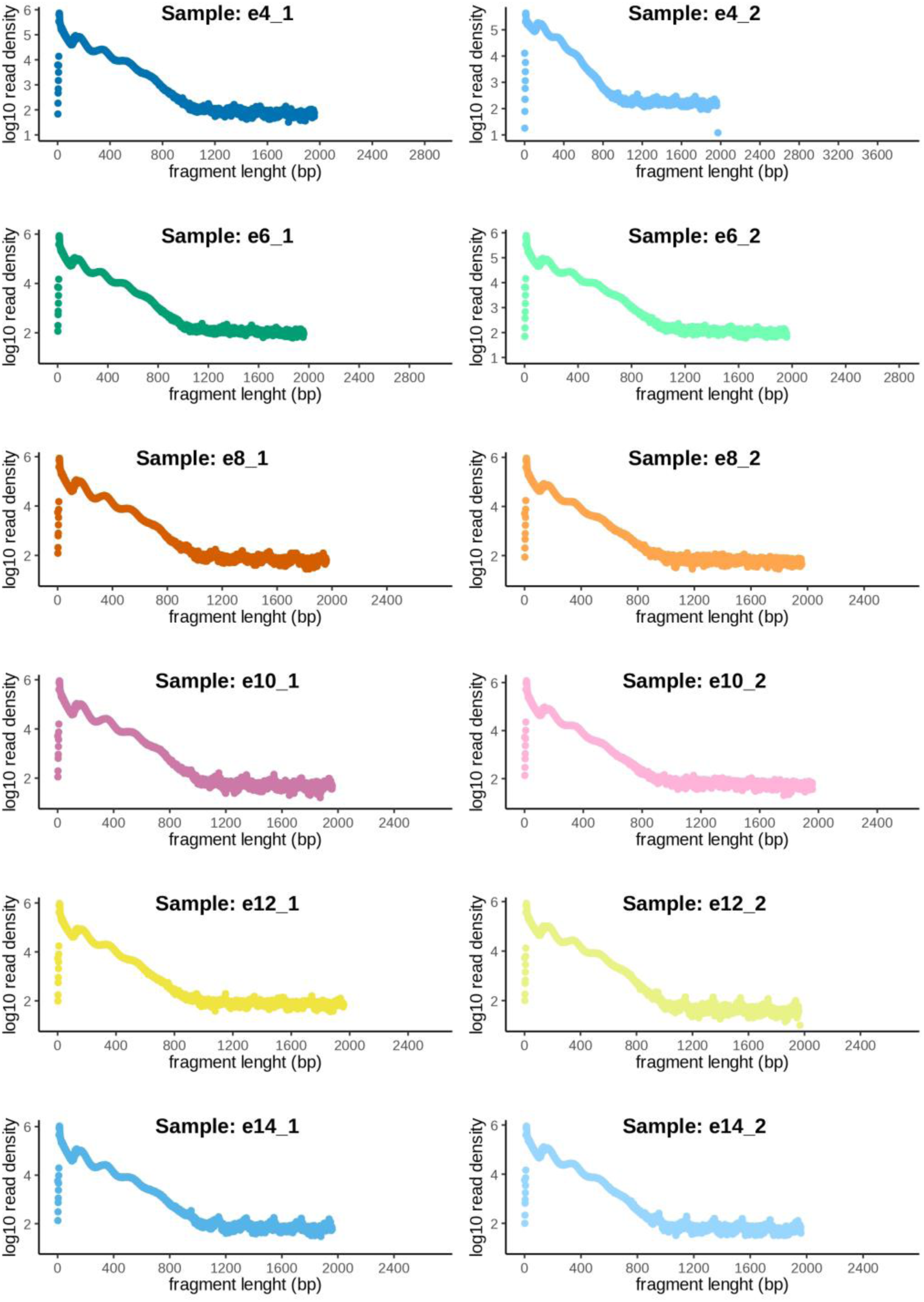
Read size distribution in ATAC-seq libraries.

**Figure S2.**
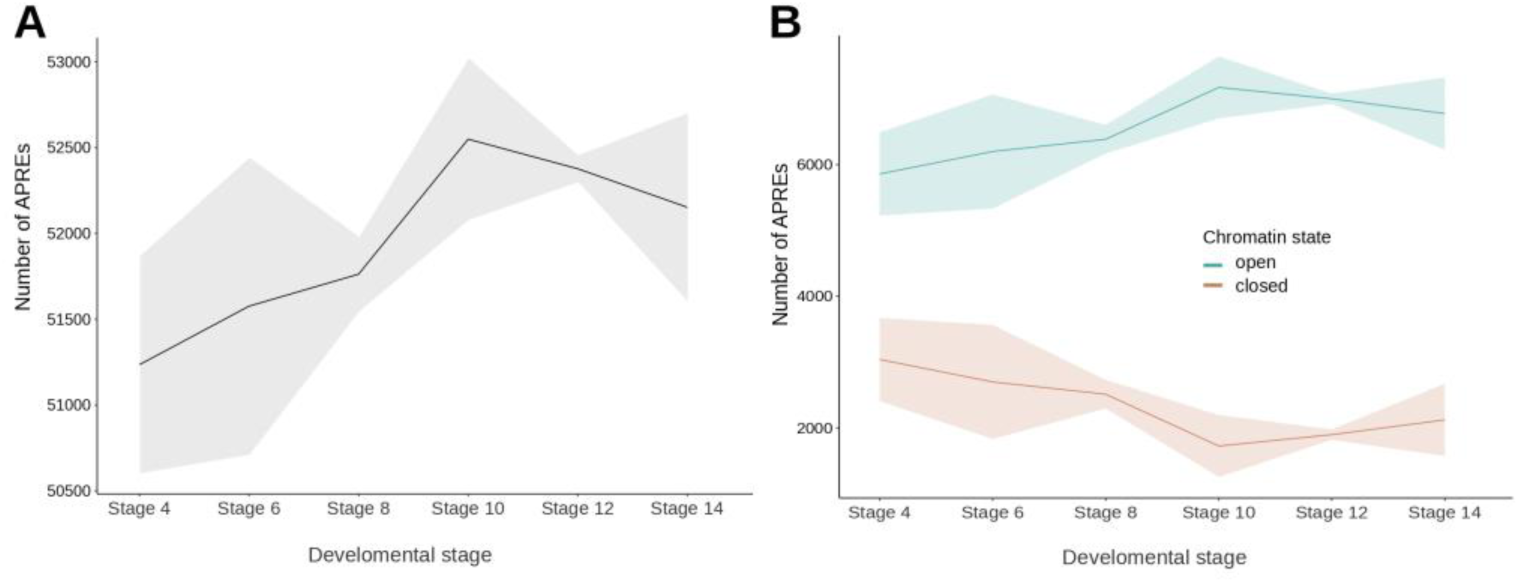
Chromatin state across different developmental stages. **(A)** Total number of APREs identified as open at each developmental stage. **(B)** Chromatin APREs that dynamically opening and closing across the stages.

**Figure S3.**
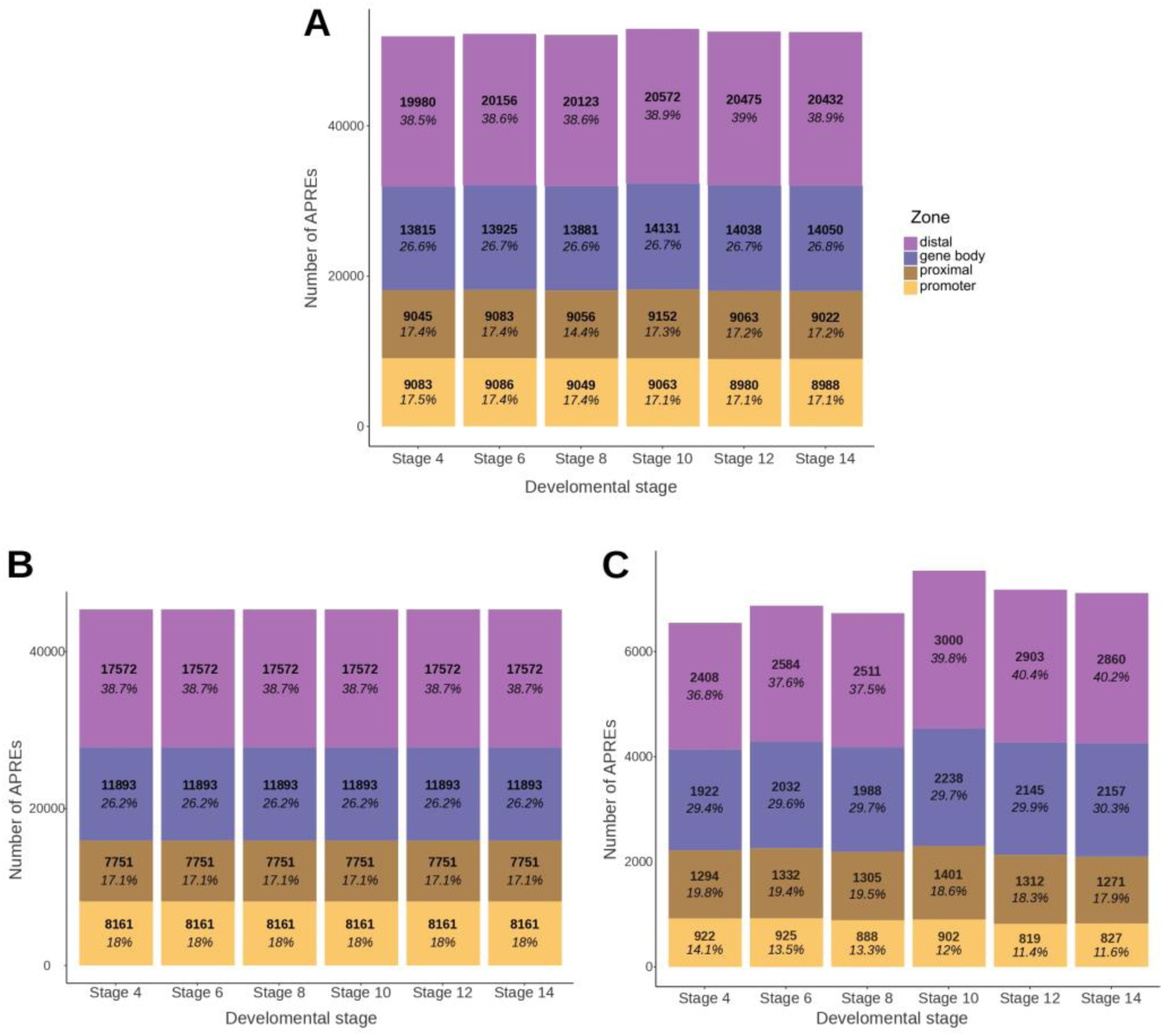
Distribution of APREs per developmental timepoint. **(A)** Number of APREs in each genomic zone distributed across each developmental stage. **(B)** Non-dynamic APREs per genomic zone distributed across each stage. **(C)** Dynamic APREs per genomic zone distributed across each stage.

**Figure S4.**
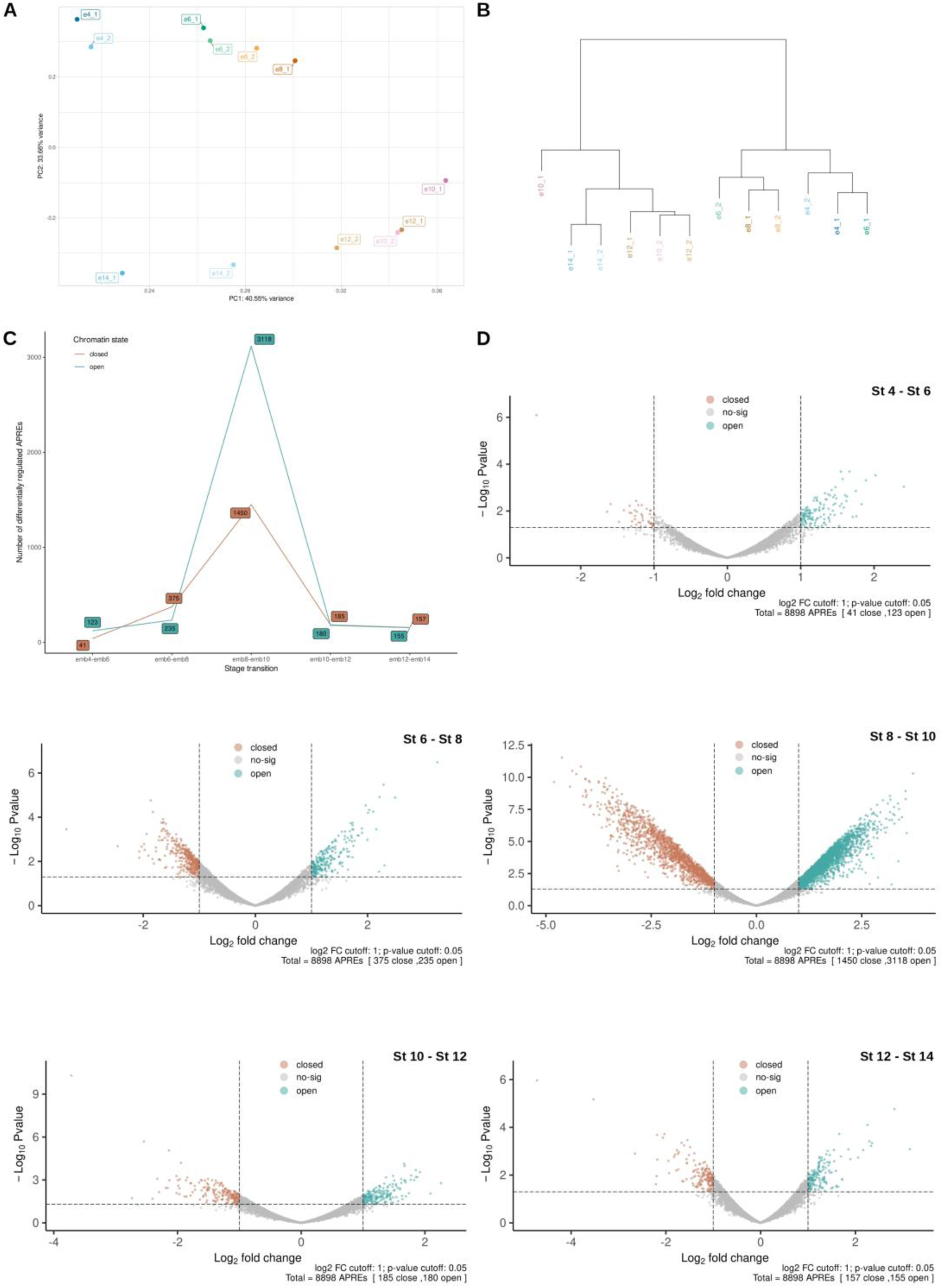
PCA analysis of ATAC libraries and differential APRE accessibility. **(A)** PCA showing the distribution of samples across the first two principal components. **(B)** Hierarchical clustering of each sample. **(C)** Number of differentially accessible chromatin regions between the different stage transitions. **(D)** Volcano plot of all differentially accessible regions for each stage transition.

**Figure S5.**
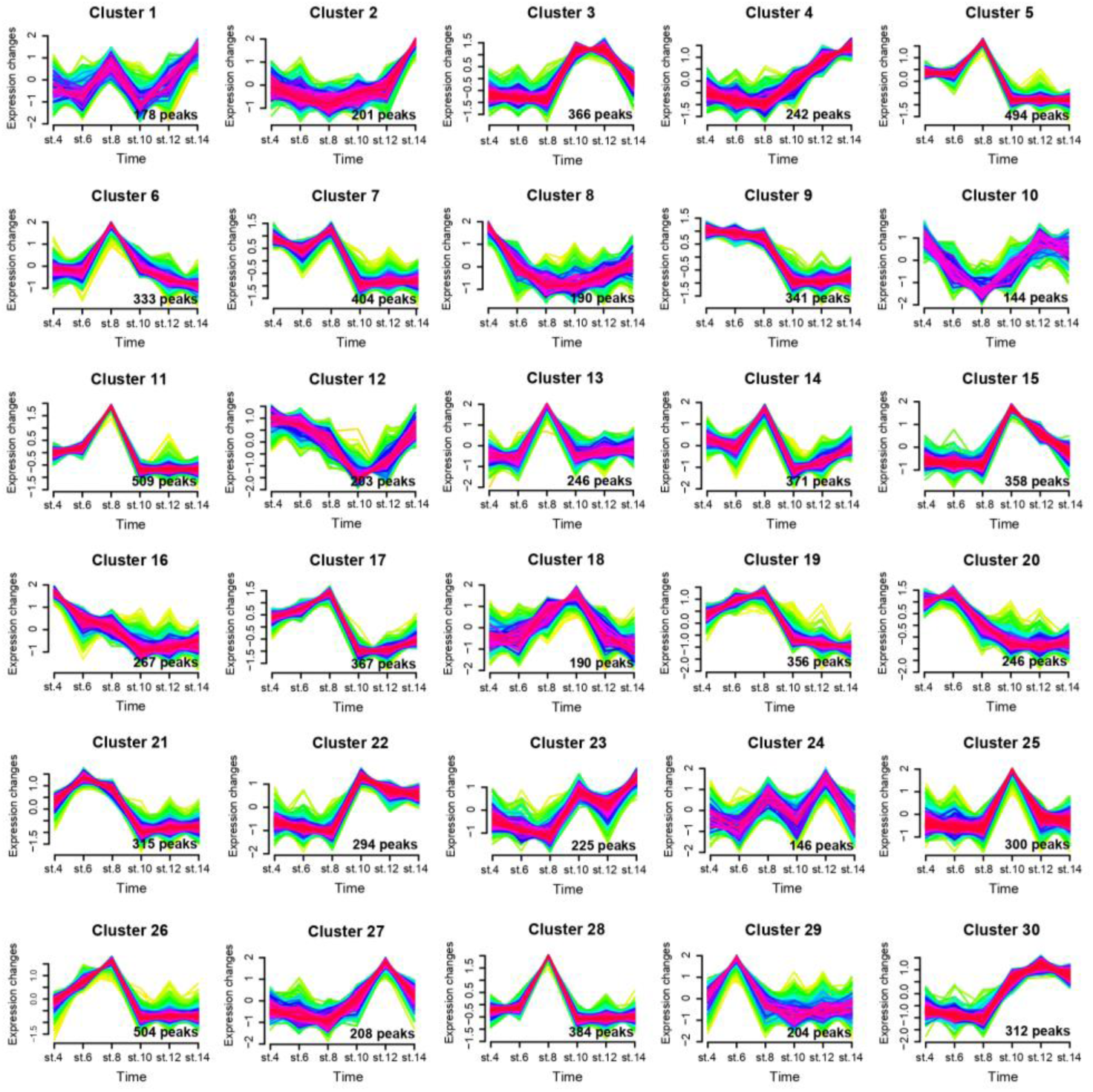
Mfuzz clustering. Patterns of chromatin accessibility across different developmental stages.

**Figure S6.**
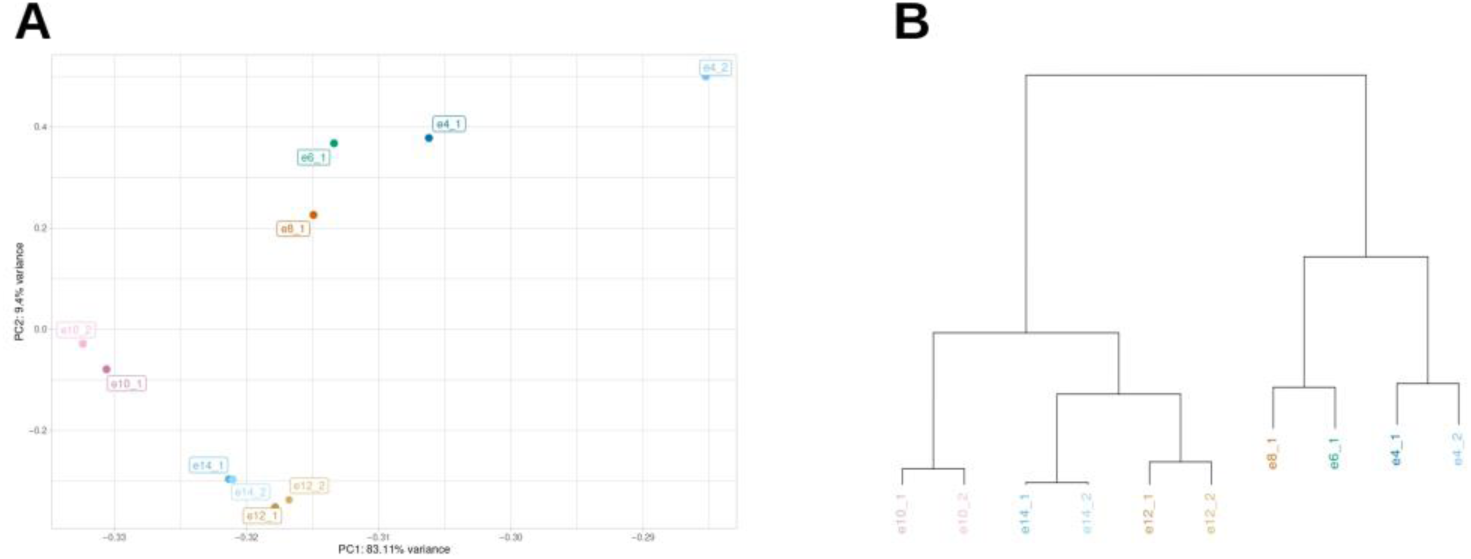
PCA analysis of RNA-seq Libraries. **(A)** PCA displaying the distribution of RNA samples across the first two principal components. **(B)** Hierarchical clustering of RNA samples.

