## Supplementary figures and images for "Gene regulatory dynamics during the development of a paleopteran insect, the mayfly *Cloeon dipterum*"

### Figure S1

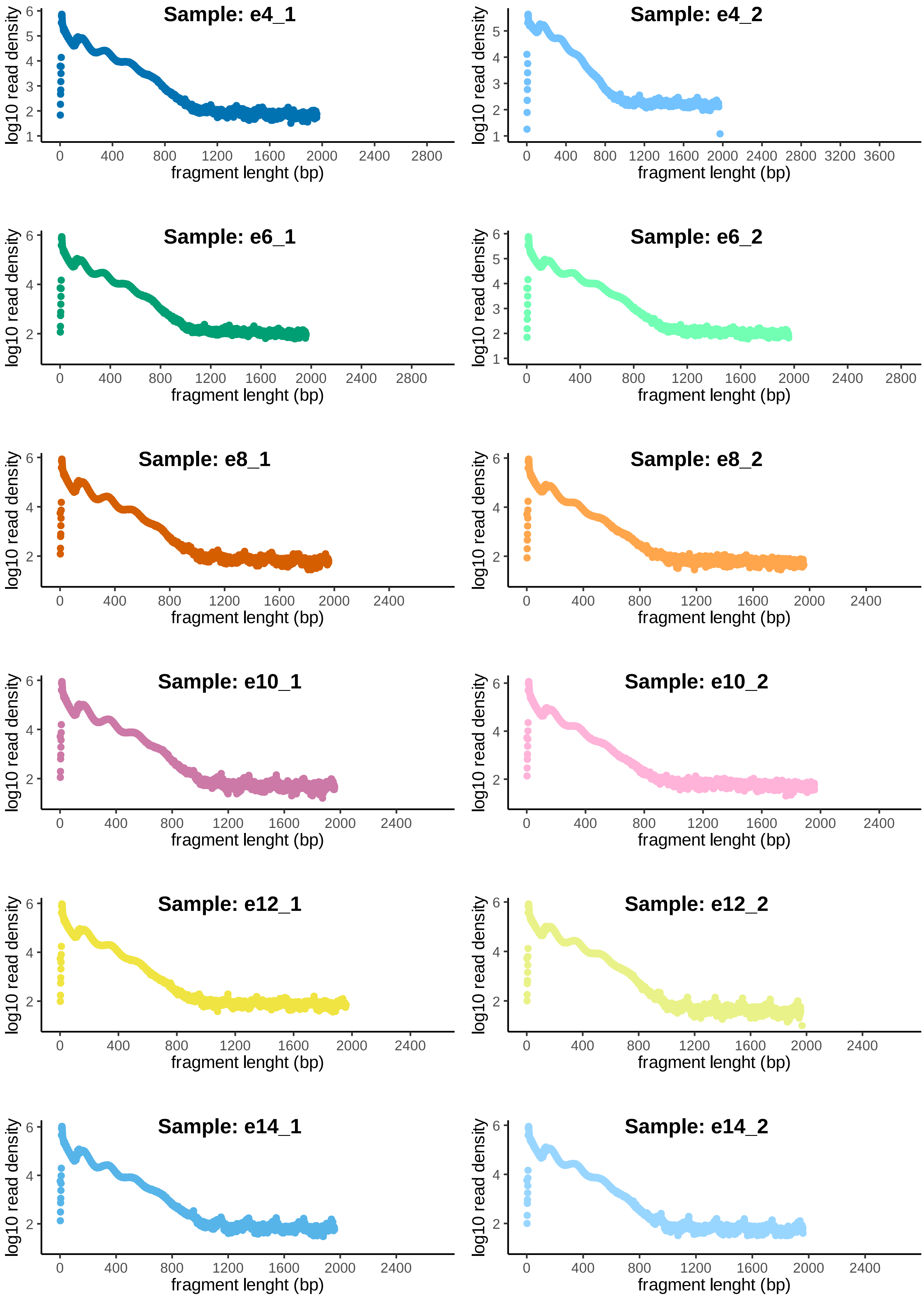

### Figure S2

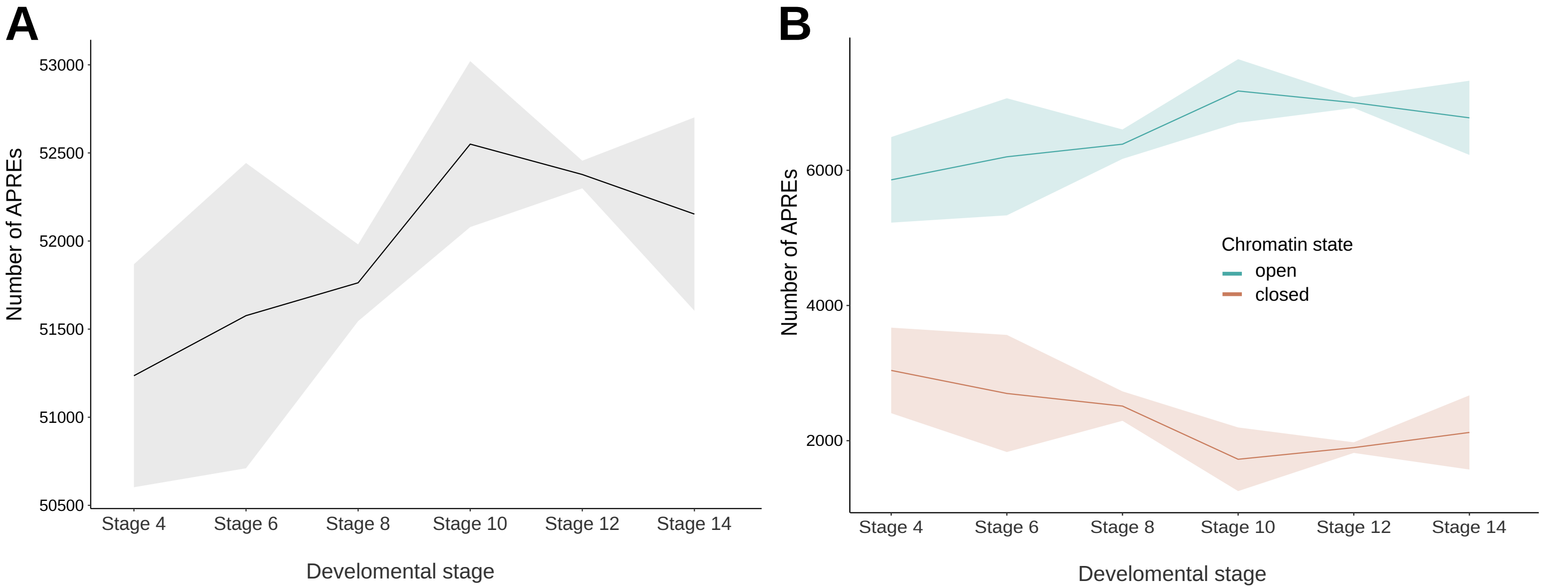

### Figure S3

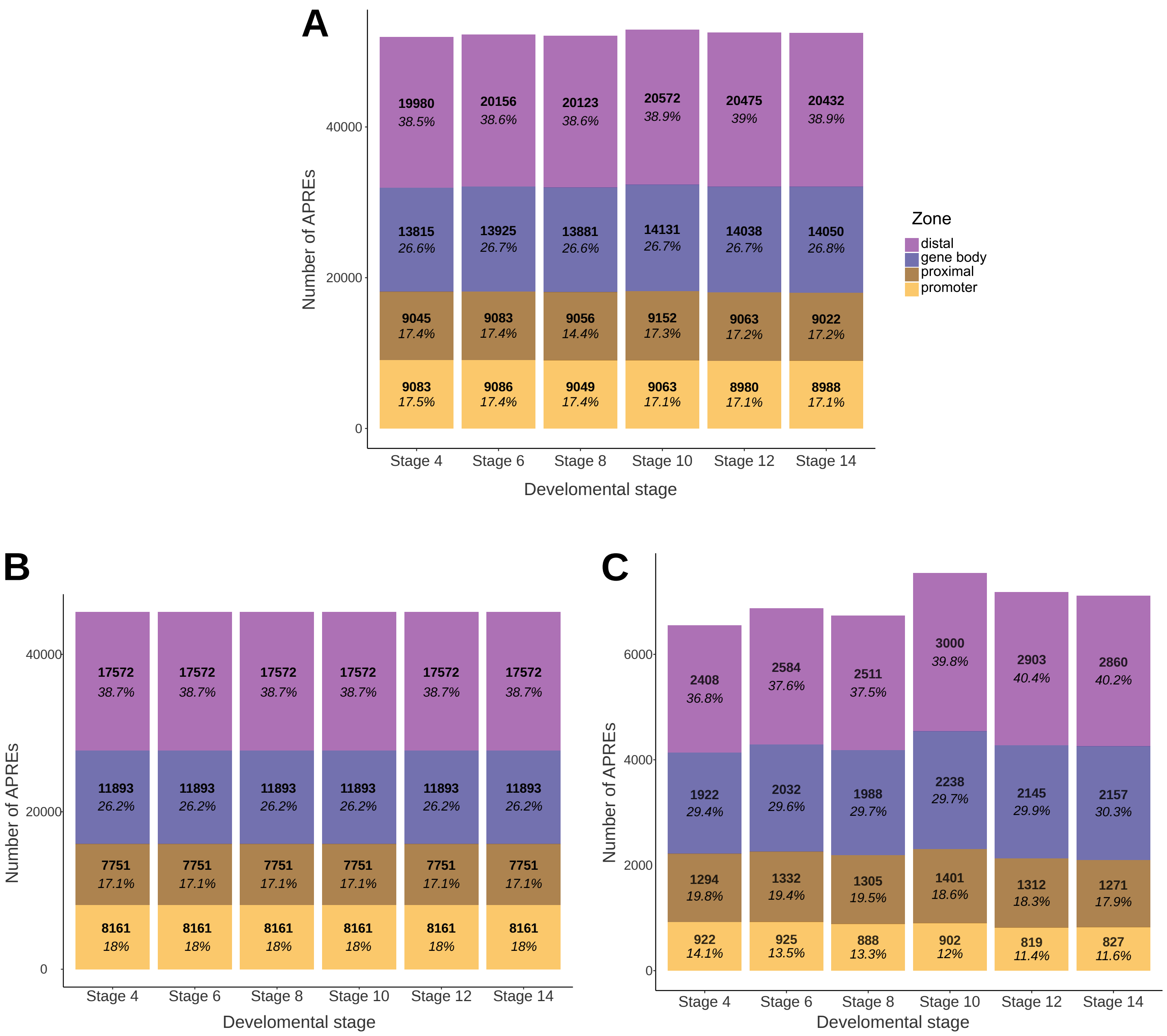

### Figure S4

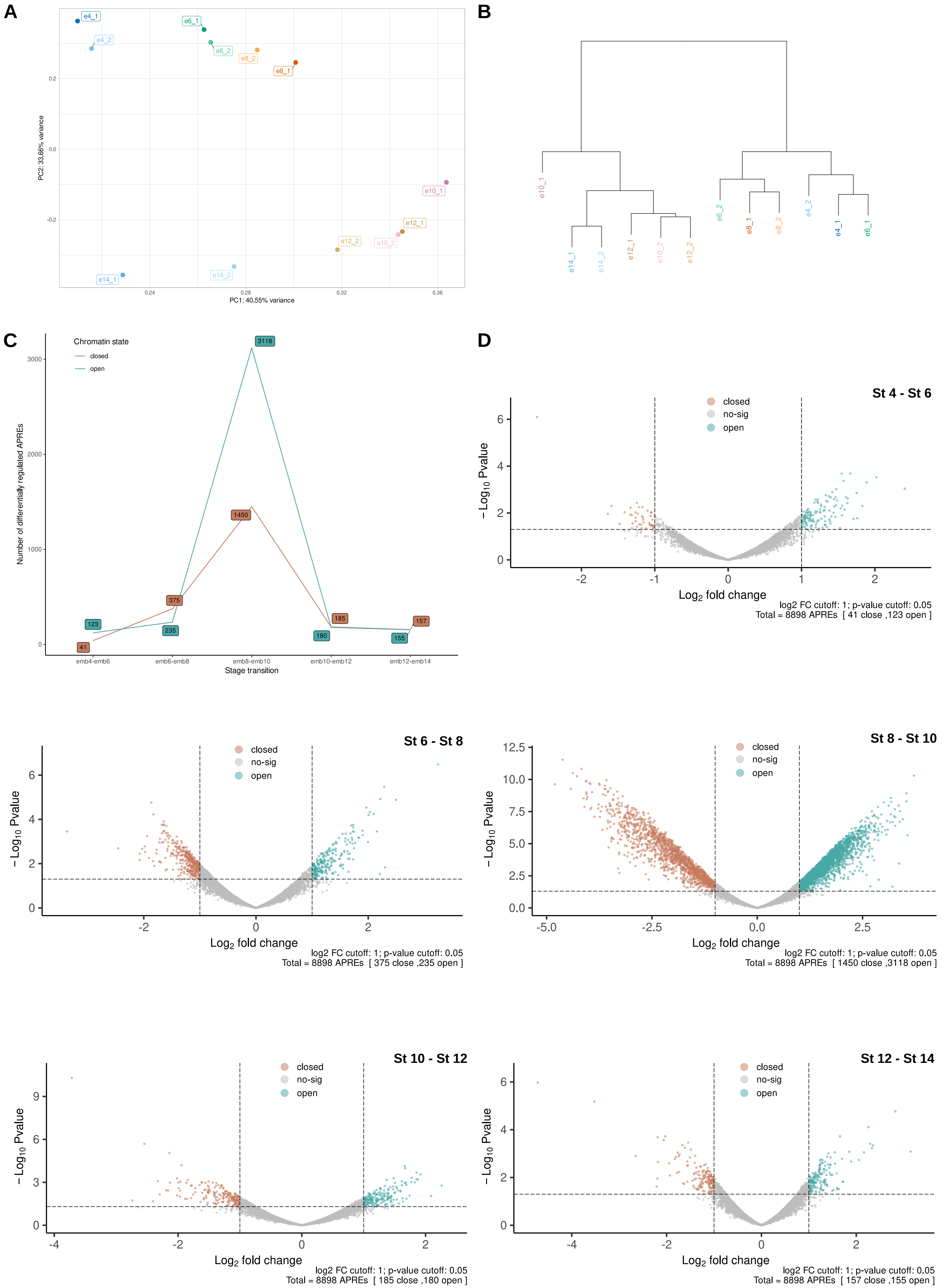

### Figure S5

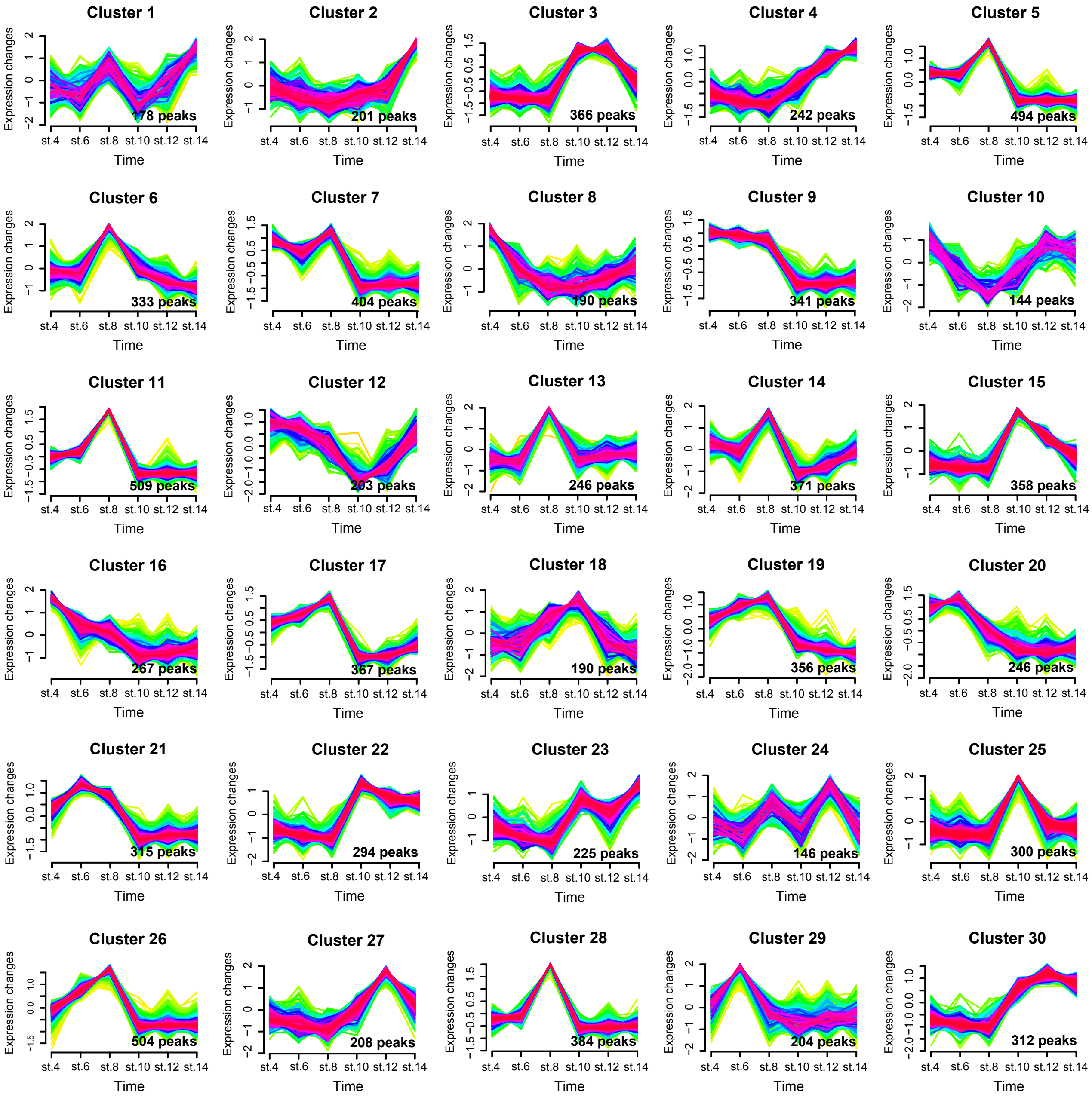

### Figure S6

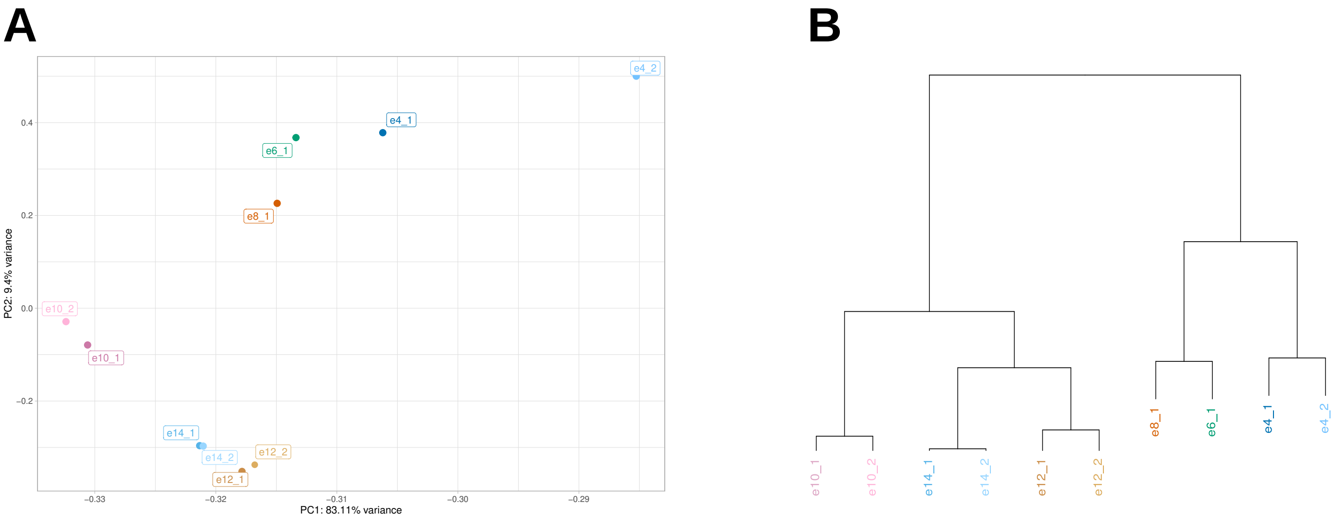
